# Signatures of Hebbian plasticity in the nanoscale morphodynamics of cortical spines

**DOI:** 10.64898/2026.08.27.747454

**Authors:** Mohammadreza Soltanipour, Aaron Nagel, Katrin I. Willig, Fred Wolf

**Affiliations:** Göttingen Campus Institute for Dynamics of Biological Networks, University of Göttingen, Göttingen, Germany; Max Planck Institute for Dynamics and Self-Organization, Göttingen, Germany; German Aerospace Center (DLR), Göttingen, Germany; Institute of Theoretical Medicine, University of Augsburg, Augsburg, Germany; Bernstein Center for Computational Neuroscience, Göttingen, Germany; Max Planck Institute for Multidisciplinary Sciences, Göttingen, Germany

## Abstract

The size, shape, and nanoscale organization of synaptic spines are predictive of physiological synaptic strength and thus key to connectome-based neural circuit models. In the intact brain, spine size and shape undergo continuous morphological remodeling. While these nanoscale morphodynamics are not well understood, it is clear that morphological changes that strongly impact synaptic strength should modify spine head and neck in a coordinated manner and that such coordinated remodeling can be triggered by spike-timing-dependent synaptic plasticity. Here we demonstrate unambiguous signatures of such coordinated spine remodeling from in vivo long-term nanoscopy of cortical spines. We infer data-driven generative models of synaptic spine morphodynamics. These models imply that although coordinated remodeling represents the smallest component of ongoing morphodynamic fluctuations, this component causes massive fluctuations in instantaneous synaptic strength. Despite these fluctuations, synaptic strengths exhibit a synapse-specific long-term persistent component that results from spine fluctuations exploring only a limited subregion of accessible spine morphospace. Our results suggest that nanoscale morphodynamic spine remodeling injects substantial fluctuations of instantaneous synaptic strength into the operation of cortical circuits. Connectome-based neural circuit models thus need to consider features of spine organization beyond spine head size and should be augmented by data-driven generative models of spine morphodynamics.

## Introduction

The operation of neuronal circuits is largely determined by their connectome, the network of synaptic interactions between neurons [1]. In mature cortical circuits, the majority of excitatory synaptic inputs is mediated by persistent dendritic spines [2, 3, 4]. The physiological strengths of these synapses are subject to short-term and long-term synaptic plasticity [5, 6] and are correlated with a spine’s morphology [7, 8, 9]. Prominently, the size of the spine head is a well established proxy of physiological synaptic strength [7]. In addition, the length and width of the spine neck also covary with synaptic strength, such that short and thick spine necks are associated with strong synapses and long and thin spine necks with weak ones [8, 9, 10]. In line with the biophysics of nanoscale neuronal compartments [10, 11, 12], the most effective way to increase synaptic strength thus involves simultaneous increases in head size and neck width coordinated with a decrease in neck length. Intriguingly, inducing spike-timing-dependent synaptic potentiation (STDP) in slices indeed causes exactly this pattern of coordinated morphological changes [7, 13, 14].

It appears natural to view the diversity of spine morphologies and synaptic strengths found in cortical circuits as the cumulative effect of changes driven by activity-dependent plasticity [15, 6]. There are, however, plausible alternatives to such a plasticity-centered view [16]. Firstly, morphological modifications might only provide a transient contribution to changes in synaptic strength with persistent modifications solely mediated by the nanoscale organization of the post-synaptic density and presynaptic active zone [17, 18, 19]. In addition, substantial components of fluctuations in spine morphology and synaptic strength might result from molecular turnover of supramolecular complexes such as a spine’s actin and tubulin cytoskeleton [20, 13, 14], its receptor clusters [18], or its postsynaptic scaffold [17]. In fact, in neuronal cultures, substantial fluctuations in spine size and molecular complement persist even in the complete absence of synaptic transmission [21, 22]. Which specific mechanisms drive the heterogeneity and dynamics of spine morphology and synaptic strength in vivo is largely unknown.

In vivo stimulated emission depletion (STED) nanoscopy offers unique possibilities to track the nanoscale morphology of individual cortical spines over periods of hours, days, and weeks [23, 24, 25]. Complementing snapshots of spine populations, provided by electron microscopic connectomics, STED nanoscopy thus opens a unique window onto ongoing synaptic dynamics [1, 26]. To address the question of persistence and volatility in spine morphodynamics, we here develop a data-driven approach to quantify, decompose and generatively model the dynamics of cortical spines in 3D morphospace. Applied to a large population of synaptic spines subserving apical tuft inputs of layer 5 pyramidal neurons, this analysis uncovers that fluctuations in spine morphology in fact exhibit signatures of STDP-induced changes and clarifies the nature of persistent and dynamic components in the dynamics of spine morphology and synaptic strength.

## Results

### Multiscale characterization reveals coordinated remodeling and a persistent baseline

Spine morphology fluctuated substantially on both short (minutes to hours) and long (days to weeks) timescales. Using two *in vivo* STED datasets spanning these regimes, we tracked head size, neck length, and neck width of individual spines (Fig. 1a–c). The absolute feature values were right-skewed; a logarithmic transform rendered their distributions approximately symmetric, whereas the changes between time points were well described by a Gaussian (Supplementary Material). In this log-transformed, *z*-scored morphospace, the short- and long-term datasets had closely matching distributions, and the averaged 68% and 95% probability contours defined a common region of morphospace (Fig. 1d).

**Figure 1:**
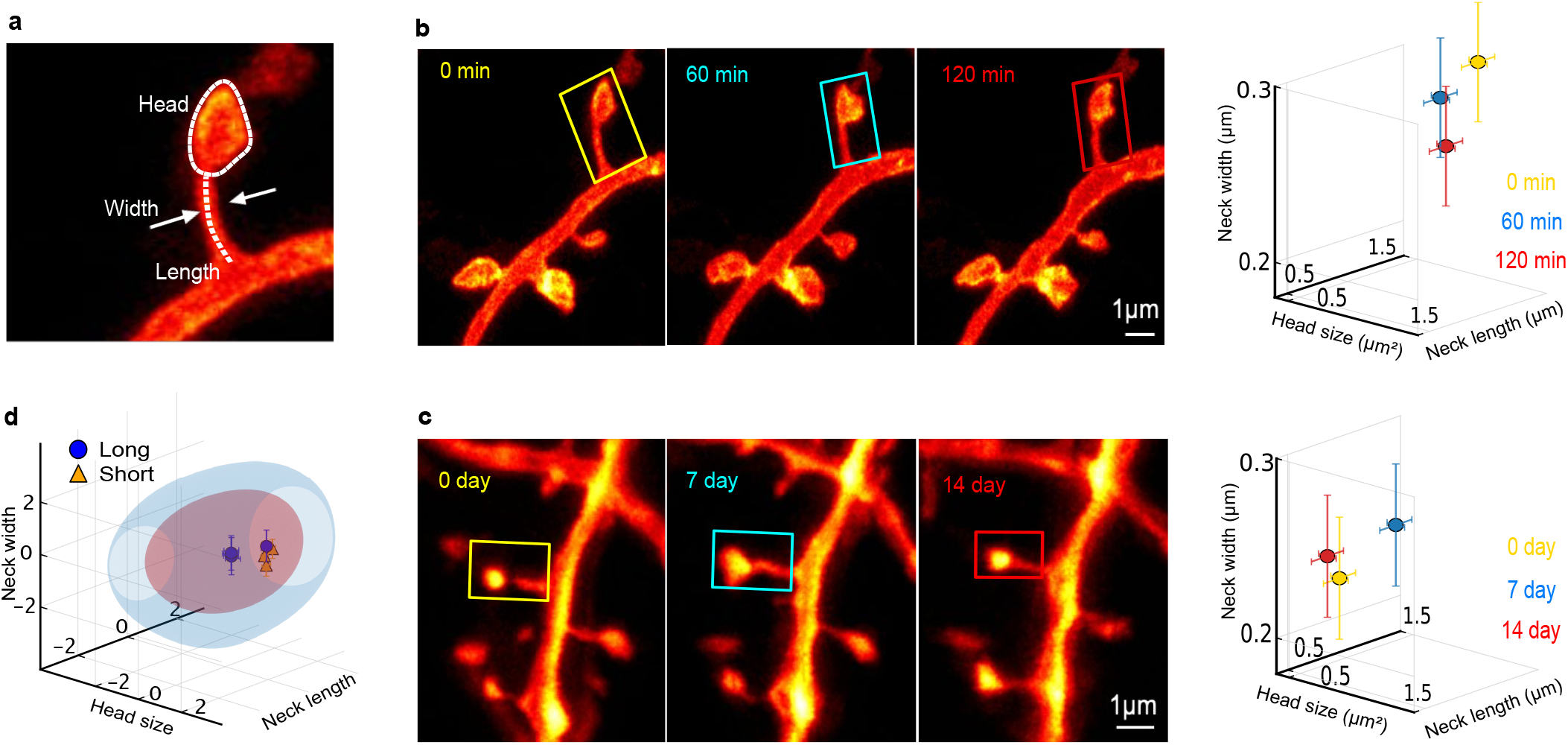
Morphological features of dendritic spines from hours to weeks. **(a)** Schematic illustrating the morphological features quantified: spine head size, neck length, and neck width. **(b, c)** Representative *in vivo* STED images and corresponding morphological measurements over short (hours, b) and long (weeks, c) time intervals. Error bars denote the estimated measurement noise for each feature, derived from the autocorrelation function at zero lag (Methods). **(d)** Averaged 68% and 95% probability contours in log-transformed, z-scored morphology space for the short- and long-term datasets.

To disentangle persistence from volatility, we computed population-averaged auto- and cross-correlation functions of the three features and inferred their parameters by Bayesian nested sampling (Figs. 2 and 3). The cross-correlations exhibited a robust sign structure—positive between head size and neck width, and negative between neck length and both (Fig. 2a). This is precisely the coordinated change—head and neck width growing while neck length shortens— that accompanies spike-timing-dependent potentiation (Fig. 2a–b), identifying a coordinated component of remodeling that carries the signature of spike-timing-dependent synaptic plasticity.

**Figure 2:**
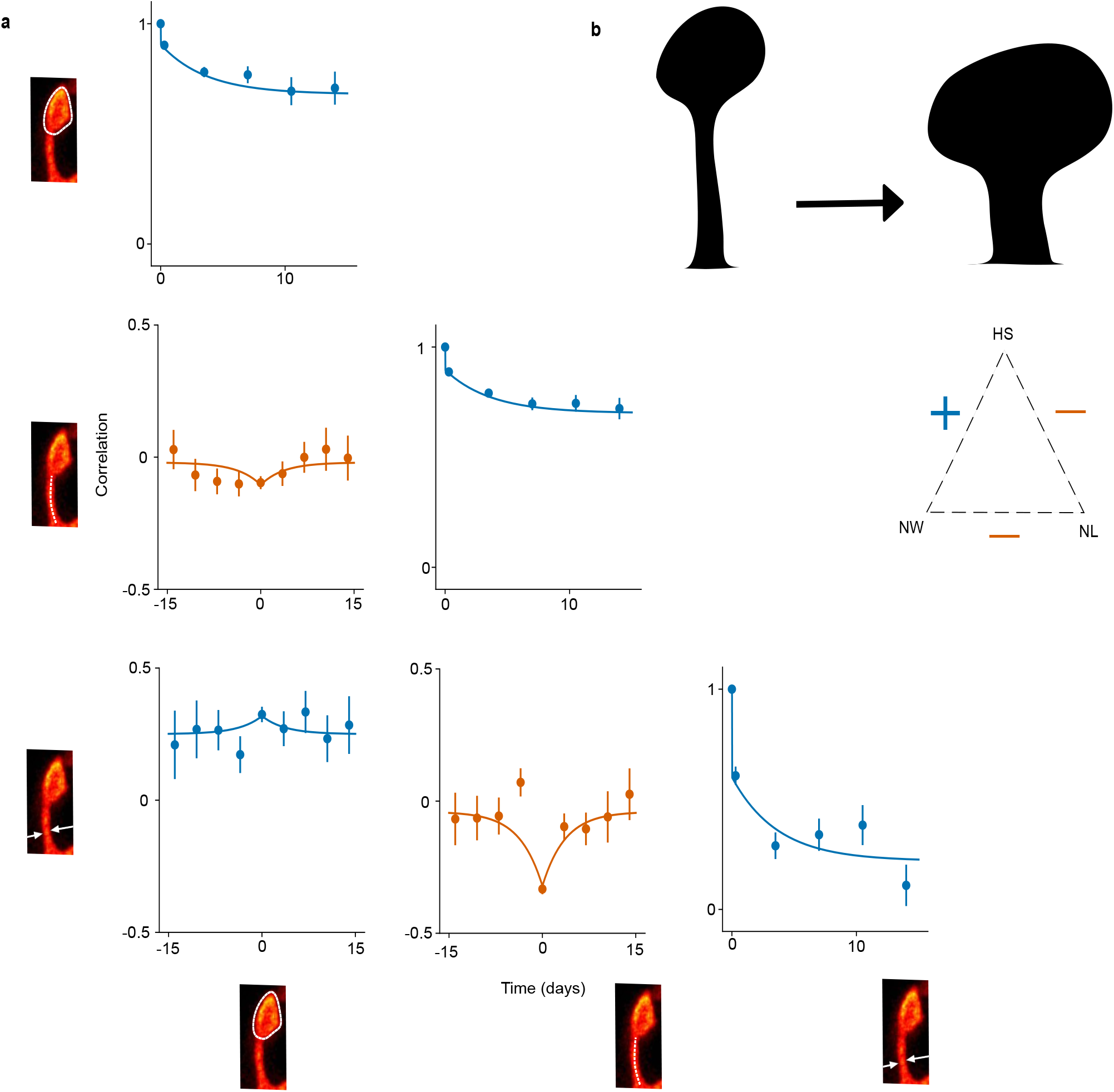
Measured temporal correlations and the expected signature of STDP-driven spine remodeling. **(a)** Measured temporal auto- and cross-correlation functions of spine head size (HS), neck length (NL), and neck width (NW). Diagonal panels show autocorrelations; off-diagonal panels show the corresponding pairwise cross-correlations. Points denote bootstrap means ± standard error; solid lines are single-exponential-plus-offset fits (Methods). Insets show representative STED images of the measured feature. **(b)** Expected morphological change under STDP-driven remodeling (left) and the corresponding pairwise correlation signs among HS, NL, and NW (right).

**Figure 3:**
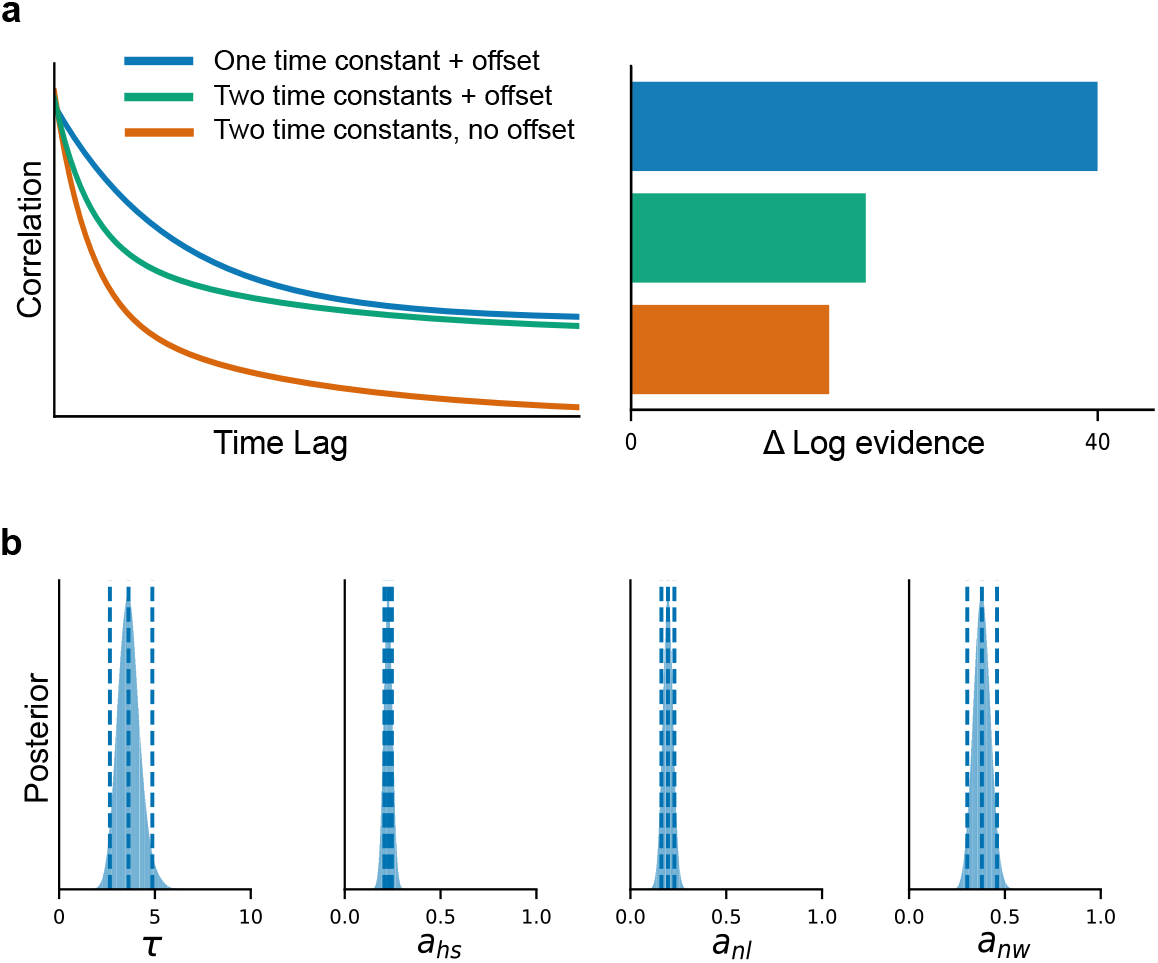
Bayesian model comparison supports a single timescale and persistent correlations in dendritic spine morphodynamics. **(a)** *Left*, schematic autocorrelation functions for the three candidate models: one time constant with offset, two time constants with offset, and two time constants without offset. *Right*, difference in log model evidence (Δ log *Z*) relative to a constant null model lacking temporal structure. The model with one time constant and a persistent offset is best supported by the data. **(b)** Marginal posterior distributions of the parameters of the best-supported model: the characteristic timescale (*τ*) and the correlation amplitudes for spine head size (*a*_*hs*_), neck length (*a*_*nl*_), and neck width (*a*_*nw*_). Dashed lines indicate the 95% credible intervals.

The correlation functions further constrained the temporal structure of these fluctuations. All auto- and cross-correlations decayed with a single characteristic timescale, *τ* ≈ 3.6 days, onto a non-vanishing offset, *C*_*ij*_(Δ*t*) = *a*_*ij*_ *e*^−|Δ*t*|*/τ*^ + *q*_*ij*_, in which the amplitude *a*_*ij*_ captures the transient, volatile component and the offset *q*_*ij*_ a persistent, spine-specific baseline. Bayesian model comparison decisively favored this single-timescale-plus-offset description over alternatives with two distinct timescales or without a persistent offset (Fig. 3). The alternative models yielded either overlapping or poorly constrained posterior distributions of timescales (Supplementary Material). The persistent offset therefore reflects genuine, quenched heterogeneity across spines rather than an artifact of the fit. Together, these results show that spine morphology combines a volatile component, decaying over a few days, with a persistent, spine-specific baseline— reconciling continuous remodeling with long-term structural identity.

### An event-based model of spine morphodynamics

To interpret the measured correlations mechanistically, we introduced a stochastic, event-based model of spine morphodynamics (Fig. 4). Each feature—head size (*h*), neck length (*l*), neck width (*w*)—is modulated by discrete, temporally sparse events drawn from a Poisson process. For spine *k* and feature *i* ∈ {*h, l, w*},

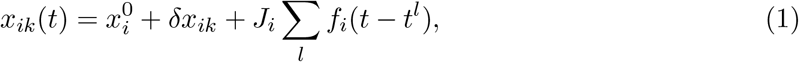

where 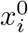 is the population mean, *δx*_*ik*_ a constant, spine-specific quenched offset, {*t*^*l*^} the event times, and *J*_*i*_ the change per event. The kernel *f*_*i*_(*t*) sets each event’s temporal profile,

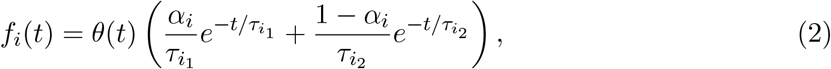

with *θ*(*t*) the Heaviside function, 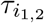 decay timescales, and *α*_*i*_ their relative weight (Fig. 4a). Each *J*_*i*_ may be drawn from a distribution spanning both potentiation and depression events; since the correlations depend on the amplitudes only through ⟨*J*_*i*_*J*_*j*_⟩, we carry a fixed *J*_*i*_ without loss of generality (Supplementary Material). The predicted covariance structure, including its sign pattern, is therefore independent of the amplitude distribution and of the direction of individual events.

**Figure 4:**
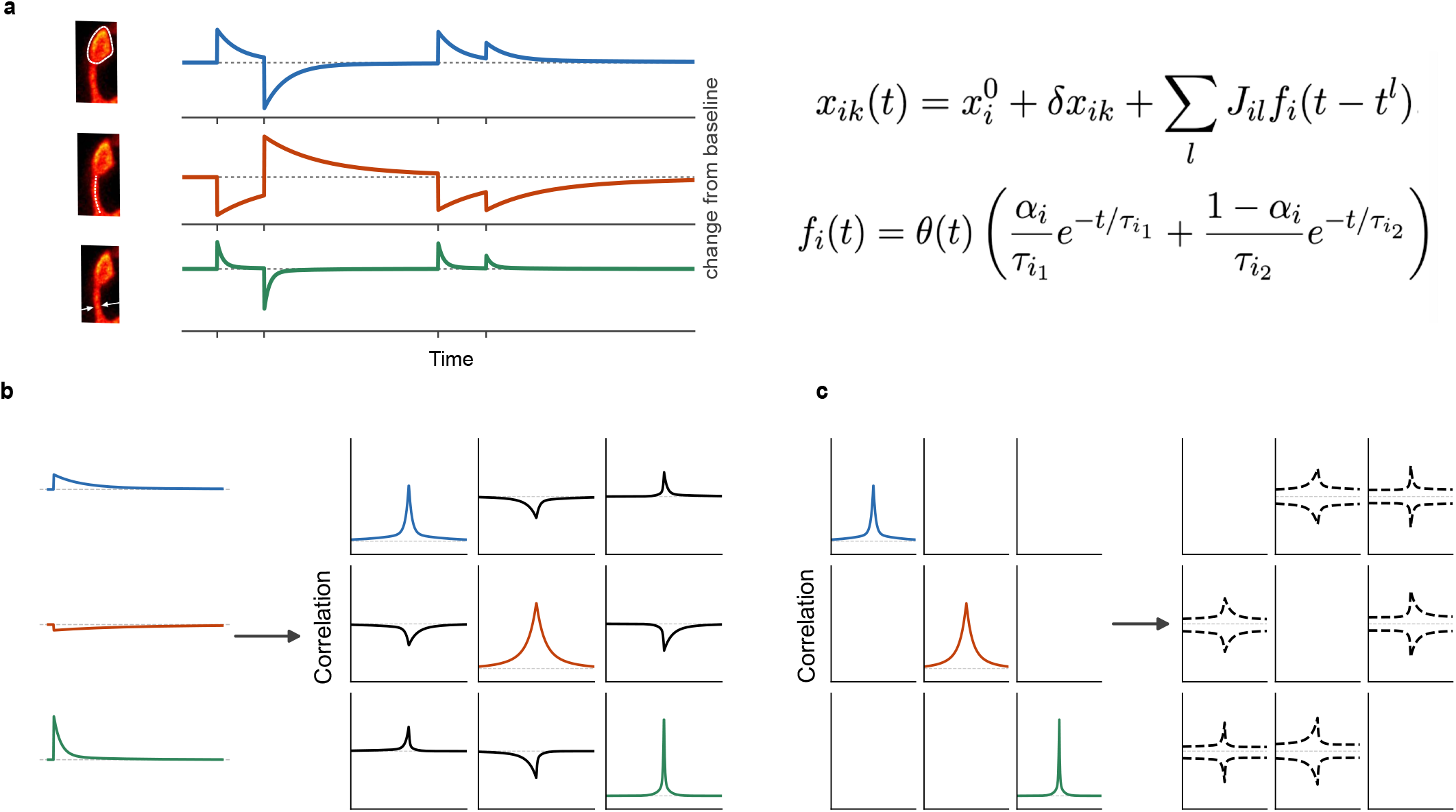
Event-based model of spine morphodynamics. **(a)** Traces (left) illustrate representative event-driven fluctuations of each feature about its baseline. Each feature—head size (blue), neck length (orange), neck width (green)—changes according to a plasticity kernel *f*_*i*_, here a sum of two decaying exponentials (timescales *τ*_*i*1_, *τ*_*i*2_; mixing weight *α*_*i*_; equations, right). The jump amplitudes *J*_*i*_ are drawn from a distribution that captures both potentiation and depression events. **(b)** Forward direction: given the kernels, the model yields the full auto- and cross-correlation matrix (3 × 3; diagonal, auto-correlations HS, NL, NW; off-diagonal, pairwise cross-correlations). **(c)** Inverse direction: from the measured autocorrelations the kernels are inferred, and the cross-covariances are then predicted from these inferred kernels (dashed).

The kernels determine the full auto- and cross-correlation matrix. For features *i* and *j*,

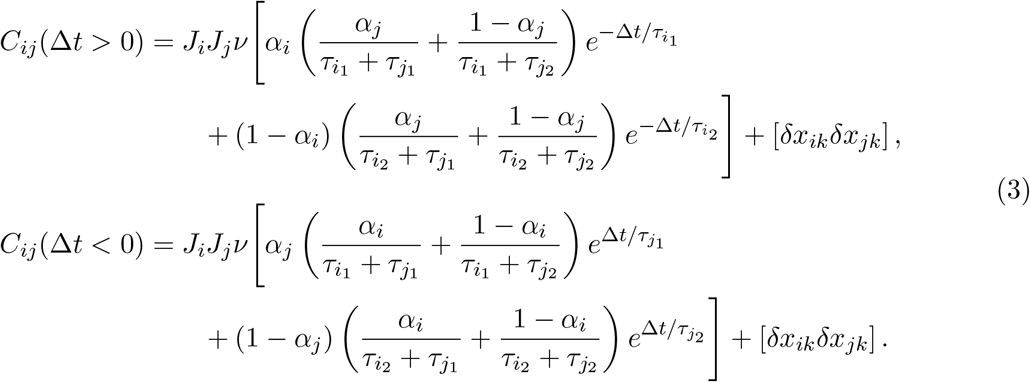

Consistent with the single timescale identified above (Figs. 2 and 3), we set *α*_*i*_ = 1 and 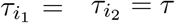, reducing the correlations to a single exponential onto a quenched offset,

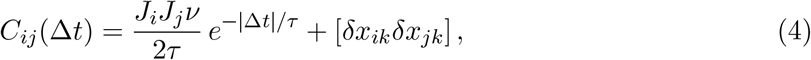

matching both the decaying transient and the non-vanishing asymptote of the data. A purely activity-independent model, by contrast, predicts only fast decay and vanishing cross-correlations (Supplementary Material).

In the forward direction, a given set of kernels fixes the complete correlation matrix (Fig. 4b). We exploit its inverse: from the measured autocorrelations we infer each feature’s kernel, and from these predict the cross-covariances (Fig. 4c). Because the same events drive multiple features, this yields a parameter-free prediction—the cross-covariance of any two features is set by their auto-covariances—and thus a direct test of whether a single common driver underlies the observed morphodynamics. To allow distinct event types (e.g. head-only events), we extend the model to several event classes, generalizing eq. (4) to

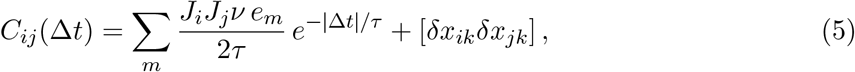

where *e*_*m*_ is the fraction of events of class *m*, with ∑_*m*_ *e*_*m*_ = 1.

### Head–neck covariance rejects a single-driver model

A single common driver makes a sharp prediction: the cross-covariance between any two features follows directly from their auto-covariances. For the head–neck covariances, this prediction falls well outside the 95% confidence intervals of the data (Fig. 5d), ruling out the simplest model in which all features are governed by one sequence of events with fixed relative amplitudes.

**Figure 5:**
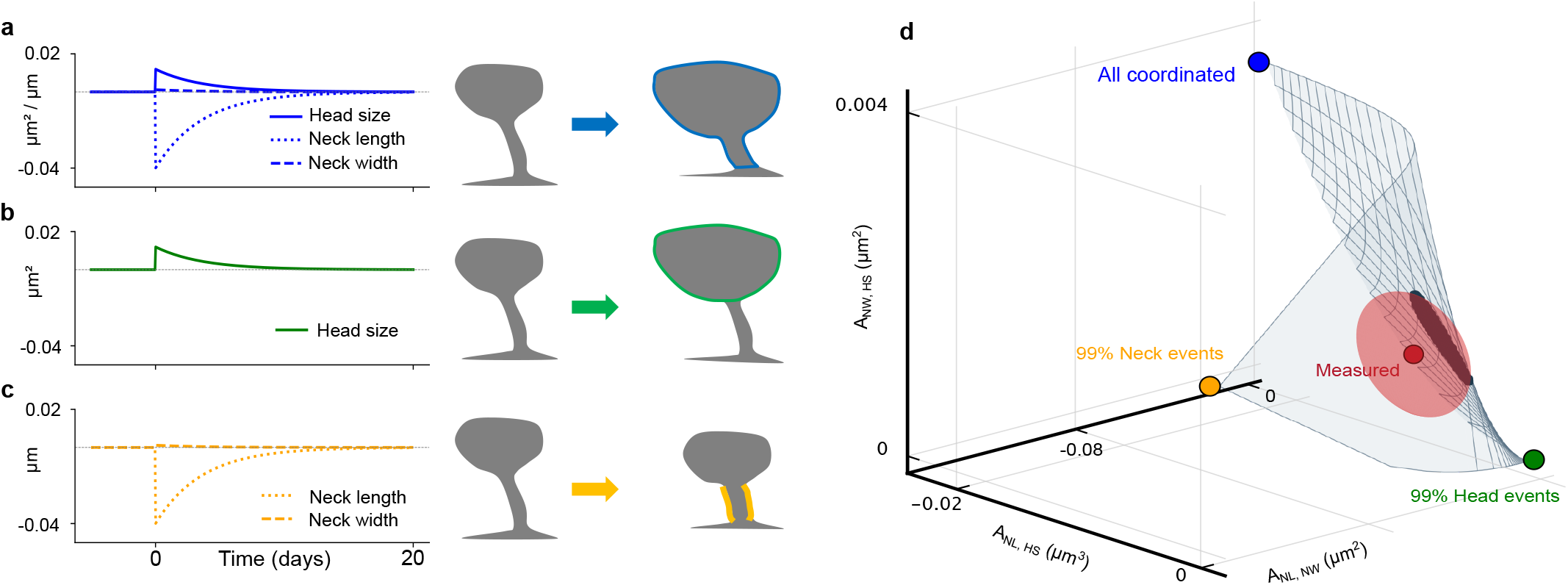
Covariance structure indicates three distinct event classes. **(a–c)** Inferred plasticity kernels describing the temporal profiles of the three event classes— coordinated events (a), head-only events (b), and neck-only events (c)—shown together with the corresponding schematic morphological change. **(d)** Model predictions for the cross-covariance amplitudes generated by different mixtures of coordinated, head-only, and neck-only events. Colored spheres denote the limiting cases of purely coordinated (blue), purely head-only (green), and purely neck-only (yellow) events; the shaded surface spans all possible mixtures. The experimentally measured covariance amplitudes (red) lie on the predicted surface, constraining coordinated events to 0.12–0.27, head-only events to 0.34–0.47, and neck-only events to 0.32–0.48 of the total, demonstrating that the observed spine dynamics are quantitatively explained by a combination of coordinated and feature-specific remodeling events.

We therefore considered an extended model with three event classes —coordinated, head-only, and neck-only—whose kernels we inferred from the autocorrelation functions (Fig. 5a–c). No single class reproduces the measured covariances. When all three are allowed, however, the manifold of achievable covariance amplitudes intersects the experimental values (Fig. 5d), constraining coordinated events to *e* = 0.12–0.27, head-only events to *e*_1_ = 0.34–0.47, and neck-only events to *e*_2_ = 0.32–0.48 of the total. All three classes thus contribute significantly, establishing that spine volatility arises from a mixture of coordinated and feature-specific remodeling.

### Coordinated events generate broad synaptic-strength fluctuations

Finally, we cast the inferred covariance structure into a generative model that reconstructs full morphological trajectories from sparse measurements (Fig. 6a; Methods). Sampling stochastic trajectories consistent with the measured auto- and cross-covariances, the model interpolated realistic fluctuations between the measured time points and reproduced both the transient and persistent components of the correlation structure. Because each trajectory draws a new realization of the volatile component while the quenched offsets are held fixed, the persistent component remained stable across the ensemble, confirming its robustness to ongoing remodeling.

**Figure 6:**
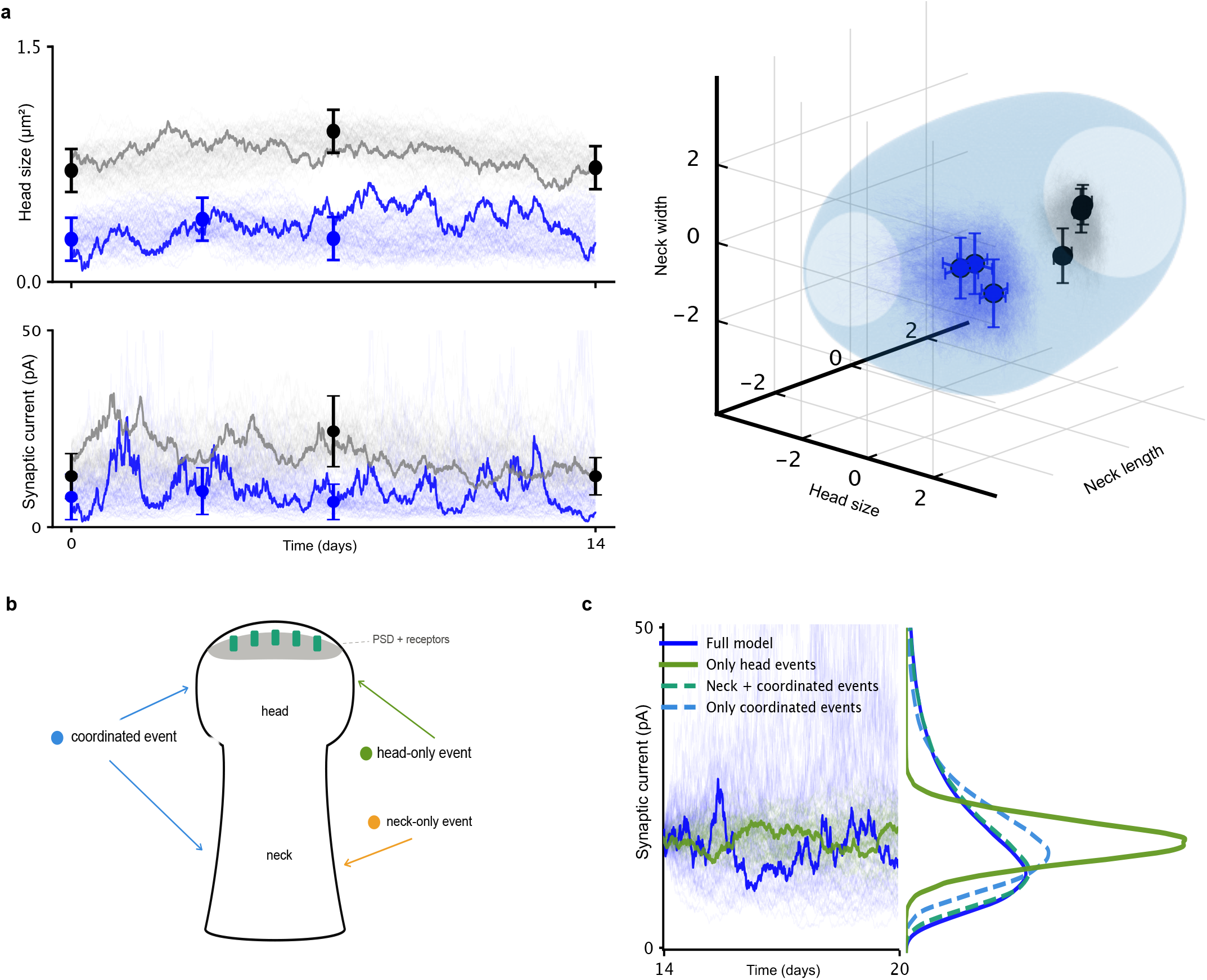
Generative reconstruction of spine morphology and synaptic current trajectories. **(a)** Reconstructed trajectories of spine head size and the corresponding synaptic current for representative large (black) and small (blue) spines (left), and the same reconstructed morphology trajectories projected into the population log-transformed, *z*-scored morphology space with the 95% probability contour (right). Filled circles denote experimental measurements with estimated measurement noise. **(b)** Schematic of the three event classes—coordinated, head-only, and neck-only (cf. Fig. 5)— acting on a single spine. **(c)** Predicted synaptic current trajectories for a representative small spine over days 14–20. The full model (blue) is compared with three counterfactual reconstructions that retain only subsets of the event classes defined in Fig. 5: only head-only events (orange), neck-only plus coordinated events (blue dashed), or only coordinated events (green dashed). Adjacent curves show the corresponding distributions of synaptic current.

To relate morphology to function, we expressed instantaneous synaptic strength as *S* ∝ *h w*^*γ*^*/l*, increasing with head size and neck width and decreasing with neck length. As spine-neck resistance is non-ohmic and departs from the naive *R* ∝ *L/r*^2^ scaling [12], we set *γ* = 1 (Methods). Inferred along each reconstructed trajectory (Fig. 6a), the synaptic current fluctuated over a broad range. Although the underlying morphological changes are Gaussian, the multiplicative form of *S* renders the resulting current distribution strongly right-skewed (Fig. 6c).

Counterfactual reconstructions isolated the origin of this variability (Fig. 6b,c). Retaining only head-only events narrowed the current distribution to a symmetric, Gaussian range, whereas retaining only coordinated events preserved a broad, right-skewed distribution close to that of the full model (Fig. 6c). Thus the small fraction of coordinated events drives the broad distribution of instantaneous synaptic strength, linking STDP-like coordinated remodeling to the dynamic range of synaptic efficacy.

## Discussion

### Summary of findings

In this study, we combined longitudinal *in vivo* STED nanoscopy with a data-driven modeling framework to quantify, decompose, and generatively model the nanoscale morphodynamics of cortical spines. Population-averaged auto- and cross-correlation functions of head size, neck length, and neck width are well described by a single-exponential decay (*τ* ≈ 3.6 days) superimposed on a non-vanishing, spine-specific quenched offset (Figs. 2 and 3). Notably, the cross-correlations exhibit a consistent sign structure—positive for head size and neck width, negative for neck length against both—exactly the coordinated pattern expected from STDP-induced potentiation (Fig. 2). An event-based model reveals that no single common driver can account for the measured head–neck covariances; instead, the data require a mixture of coordinated and feature-specific events, with coordinated events constituting only 12–27% of the total (Fig. 5). Although this coordinated component is the smallest fraction of ongoing morphodynamics, it accounts for a dominant share of the predicted fluctuations in instantaneous synaptic strength (Fig. 6). These fluctuations, however, remain confined to a limited subregion of morphospace centered on a synapse-specific mean, so that synaptic strength retains a persistent, quenched component that is shielded from ongoing remodeling.

Our results provide strong evidence that both coordinated and independent components of spine morphodynamic fluctuations should be considered strictly transient [16]. Although the coordinated component of these fluctuations is most parsimoniously explained by STDP-driven processes, our results render it unlikely that the instantaneous geometry of spines encodes a persistent memory trace of past activity-dependent changes. This does not imply that the respective synapses cannot preserve such a memory. In fact, the apical tuft synapses examined by us are directly implicated in learning processes, impacting top-down modulation of cortical processing [27], and the minor fraction of potentially STDP-driven coordinated changes notably accounts for a dominant fraction of changes in synaptic strength. The finite correlation time of ongoing fluctuations, however, supports a multilevel picture of synaptic plasticity, in which morphological changes transiently modify synaptic strength over a period of up to a few days, while lasting memory traces are ultimately laid down in the molecular organization of the synapse such as in the patterning of PSD subclusters [18, 17] or trans-synaptic columns aligning presynaptic release sites and postsynaptic receptor clusters [19]. Supporting this notion, PSD size and spine head size are only partially correlated and exhibit a larger degree of independence under conditions of environmental enrichment, which enhances learning processes [25].

Nanoscale morphodynamic fluctuations, although temporally transient, are predicted to cause large changes in synaptic strength and thus to mediate a substantial component of ongoing changes of synaptic strength in vivo. In morphospace, the corresponding dynamic fluctuations are centered on a synapse-specific mean and confined to a limited subregion. As a consequence, spine geometry generates a major contribution to the total heterogeneity of synaptic strengths that is shielded from modification by learning processes. While the function of such a quenched component of synaptic strengths is presently unknown, models of neural circuit function suggest several possibilities: Firstly, progress in training recurrent neural networks has uncovered that random quenched heterogeneity in connectivity can facilitate training networks that can flexibly switch between different computations [28, 29]. In terms of storage capacity, theories of recurrent cortical circuits require broad distributions of synaptic strength for optimal capacity [30]. A substantial degree of quenched synaptic heterogeneity might thus prime cortical circuits to operate near optimal capacity, if storage can be achieved by superimposed learning processes of limited dynamic range.

## Methods

### Spine morphology data

For this study, no new experimental data were generated. The long-term spine morphological changes were extracted from the supplementary material of control mice in Steffens et al. [24]. For short-term changes, we used the super-resolution STED imaging of control mice in Wegner et al. [25], where spine heads were originally quantified. Here, we additionally assessed spine neck length and width by manual measurements, as performed for the long-term measurements in Steffens et al. [24]. We performed the additional analysis on the same spines as the published head sizes in Wegner et al. [25]. Spine neck length was measured by drawing a line along the neck from the dendrite to the beginning of the spine head using the freehand line tool in Fiji [31]. Spine neck width was measured as the FWHM of a line profile at the location of the smallest neck width. Thus, both datasets were analyzed in the same way. The labeling methods, however, differed slightly: a volume expression of EGFP in a transgenic mouse line was used for the long-term changes in Steffens et al. [24], whereas an EGFP membrane labeling by viral expression vectors was used for the short-term measurements in Wegner et al. [25]. The long-term changes were measured in motor cortex, while the short-term changes were measured in visual cortex. Both studies, however, examined spine morphology in layer 1 apical dendrites of pyramidal neurons.

The number of spine measurement pairs for each time shift is shown in Table S1, which were used for scatter plots and autocorrelation calculations. The number of data points used for the corresponding cross-correlations, combining data pairs of different features at specific time shifts, is shown in Table S2.

### Modeling and analysis

For each morphological feature, we computed auto-correlations by pooling all same-spine measurement pairs separated by a given time lag. Cross-correlations were calculated analogously across different features at matched time points. Pearson correlation coefficients were used throughout. To quantify uncertainty, we applied nonparametric bootstrap resampling (*n* = 100) of the paired observations and report the standard error across resamples. The covariance matrix was obtained as *C*_*ij*_(Δ*t*) = *r*_*ij*_(Δ*t*) *σ*_*i*_*σ*_*j*_, where *r*_*ij*_(Δ*t*) is the correlation coefficient at lag Δ*t* and *σ*_*i*_, *σ*_*j*_ are the empirical standard deviations of features *i* and *j*.

### Bayesian inference and data-driven generative model

To characterize the temporal structure of morphological fluctuations, we compared candidate functions (single-or double-exponential decays with and without an offset) to the empirical auto- and cross-correlation data and an additional term *n*_*i*_ accounting for the measurement noise, *C*_*ii*_(Δ*t*) = *F*_*ii*_(Δ*t*) + *n*_*i*_ *δ*_Δ*t*,0_. Cross-correlations were modeled analogously, but without the noise term, assuming noise measurements are independent, *C*_*ij*_(Δ*t*) = *F*_*ij*_(Δ*t*). Parameters were inferred using Bayesian nested sampling (Dynesty [32]), assuming Gaussian errors from bootstrap estimates of the empirical correlations. Posterior distributions yielded credible intervals for all parameters. The data were well explained with one exponential function with an offset. Two-exponential models were rejected (see Supplementary material).

To generate surrogate trajectories, we used the fact that correlation functions decay exponentially with time constant *τ* = 3.6 days, and constructed the corresponding spectral density 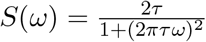. Multivariate Gaussian processes were then sampled in the frequency domain with covariance matrices matched to the empirically inferred auto- and cross-covariance amplitudes. After inverse Fourier transformation and rescaling, this yielded synthetic trajectories whose covariance structure reproduced the measured correlations.

### Synaptic strength estimation

We estimated an instantaneous synaptic current from each spine’s raw head size *h*, neck width *w*, and neck length *l* as *S* ∝ *h w*^*γ*^*/l*. The linear dependence on head size follows from the well-established proportionality between spine head size and synaptic strength [7]. The neck contributes through its resistance, which attenuates the synaptic current reaching the dendrite: strength therefore decreases with neck length and increases with neck width. A naive ohmic cable would give *R* ∝ *L/r*^2^ and hence *γ* = 2; however, spine-neck resistance is non-ohmic and scales more weakly with neck radius, closer to *r*^−1^ than *r*^−2^ [12]. We therefore adopted the milder scaling *γ* = 1. The proportionality constant sets only the overall scale and was fixed so that the mean strength across the long-term measurements corresponds to a typical current of 15 pA; the same constant was used for all trajectories and event classes. Because the morphological fluctuations are multiplicative, the resulting current distribution is right-skewed even though the underlying changes are Gaussian.

## Acknowledgments

We thank the authors of the original experimental studies that formed the basis of this work. In particular, we acknowledge Steffens et al. [24] for the long-term in vivo STED imaging data and Wegner et al. [25] for the short-term measurements. Furthermore, we thank Silvio Rizzoli, Michael Fauth, Gasper Tkacik and Matthias Häring for fruitful discussions. This work was supported by the Deutsche Forschungsgemeinschaft (DFG, German Research Foundation) 436260547 in relation to NeuroNex (National Science Foundation 2015276) and under Germany’s Excellence Strategy -EXC 2067/1-390729940, by the DFG -Project-ID 317475864 -SFB 1286 and Project-ID 430156276 -SPP 2205. This work was further supported by a Research Grant from HFSP (Ref.-No: RGP025/2023), and by the Niedersächsisches Vorab of the VolkswagenS-tiftung through ZN3420 and through the Göttingen Campus Institute for Dynamics of Biological Networks (ZN3326 & ZN3371).

## Supplementary material

**Table S1:** Number of data pairs from spine morphology dataset for different time shifts for minutes (m) and days (d).

| time shifts $\Delta t$ | 30 (m) | 60 (m) | 120 (m) | 3.5 (d) | 7 (d) | 10.5 (d) | 14 (d) |
| --- | --- | --- | --- | --- | --- | --- | --- |
| HS pairs | 417 | 285 | 189 | 428 | 280 | 158 | 100 |
| NL pairs | 221 | 154 | 72 | 442 | 289 | 164 | 102 |
| NW pairs | 211 | 142 | 68 | 275 | 190 | 111 | 73 |

**Table S2:** Number of data points from spine morphology dataset for different time shifts of the cross-correlation data for minutes (m) and days (d)

| $\Delta t$ | 0 | $\pm 30$ (m) | $\pm 60$ (m) | $\pm 120$ (m) | $\pm 3.5$ (d) | $\pm 7$ (d) | $\pm 10.5$ (d) | $\pm 14$ (d) |
| --- | --- | --- | --- | --- | --- | --- | --- | --- |
| NL-HS | 675 | 245 | 165 | 83 | 437 | 289 | 164 | 102 |
| NW-HS | 630 | 234 | 155 | 80 | 314 | 213 | 134 | 87 |
| NW-NL | 638 | 214 | 145 | 70 | 318 | 215 | 134 | 87 |

### Correlation functions model comparison: one vs. two timescales

To identify the best phenomenological description of the measured correlation functions, we considered three candidate models: a single exponential plus offset (Fig. S2), a double exponential plus offset (Fig. S3), and a double exponential without offset (Fig. S4). Using Bayesian inference (nested sampling), we found that the model evidence decisively favored the single-timescale-plus-offset model (Fig. 3), whose fits to the empirical correlations are shown in Fig. 2.

The posterior distributions explain this preference. The one-timescale model converged robustly to a well-defined *τ* ≈ 3.6 days (95% credible interval +1.22*/* − 0.96; Fig. S2). For the double exponential plus offset, *τ*_1_ and *τ*_2_ overlapped heavily, collapsing back to a single effective timescale (Fig. S3). Without an offset, the model was additionally forced to absorb the non-vanishing long-lag correlations into the second slow exponential, resulting in a poorly constrained timescale (Fig. S4). A spine-specific offset is thus an essential component of the description, and the data are best captured by a single exponential decay onto a persistent baseline.

### Log-normal feature distributions and Gaussian increments

The raw values of head size, neck length, and neck width were right-skewed and better described by a log-normal than a Gaussian distribution (Fig. S1, top). Their between-timepoint increments, by contrast, were Gaussian (Fig. S1, bottom). Distributions were spine-weighted and compared by weighted ΔAIC [33]; because the variance of the increments grows with time lag, each lag was standardized before pooling. Log-normal and Gaussian fits to the raw values were obtained in closed form from the (log-)mean and variance, while the shifted log-normal fitted to the standardized increments was obtained by weighted maximum likelihood, restarted from multiple initializations to avoid local optima in its three-parameter likelihood. Given full freedom to remain skewed, this shifted log-normal instead converged onto the standard normal, confirming that the increments are Gaussian.

### Derivation of correlation functions for the event-based model

To model the observed correlations in dendritic spine morphology, we consider a minimal stochastic formulation in which each morphological feature evolves through the superposition of discrete events. We assume that events follow a stationary Poisson process with constant rate *ν*, and that quenched offsets have zero mean across spines, [*δx*_*ik*_] = 0. For spine *k*, the morphology is described as:

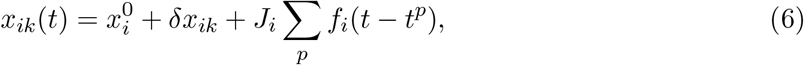

with all symbols as in the main text: *x* = (*h, l, w*) collects the three features, *p* labels events, *δx*_*ik*_ is the quenched, spine-specific offset, *J*_*i*_ the per-event amplitude, and *f*_*i*_(*t*) the plasticity kernel, taken as a normalized sum of two exponentials,

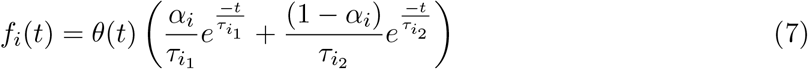

where *θ*(*x*) is the Heaviside step function and *α*_*i*_ weights the two time constants.

In the most general formulation, each event carries its own amplitude *J*_*il*_, so that 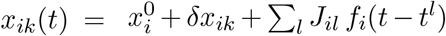, with the *J*_*il*_ drawn independently from a distribution that may span both potentiation (*J*_*il*_ *>* 0) and depression (*J*_*il*_ *<* 0) events. The correlation functions depend on these amplitudes only through the product of two events acting on features *i* and *j*; averaging over the amplitude distribution therefore replaces this product by its mean ⟨*J*_*i*_*J*_*j*_⟩. Since ⟨*J*_*i*_*J*_*j*_⟩ enters merely as an overall factor, it fixes the magnitude but not the shape or sign structure of the correlations. Replacing the random *J*_*il*_ by a single fixed amplitude *J*_*i*_ per feature, with *J*_*i*_*J*_*j*_ = ⟨*J*_*i*_*J*_*j*_⟩, thus leaves all correlation functions unchanged. We adopt this fixed-amplitude form for the remainder of the derivation without loss of generality.

The average of a spine feature reads:

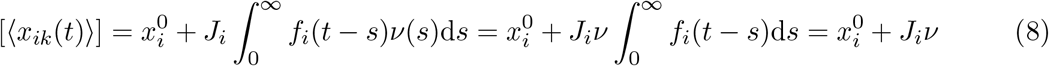

The [·] denotes the ensemble average and ⟨·⟩ the temporal average. Here, *ν* represents the plasticity rate, assumed to have reached its asymptotic value, *ν*(*t*) = *ν*.

Calculating the correlation functions is straightforward; they read:

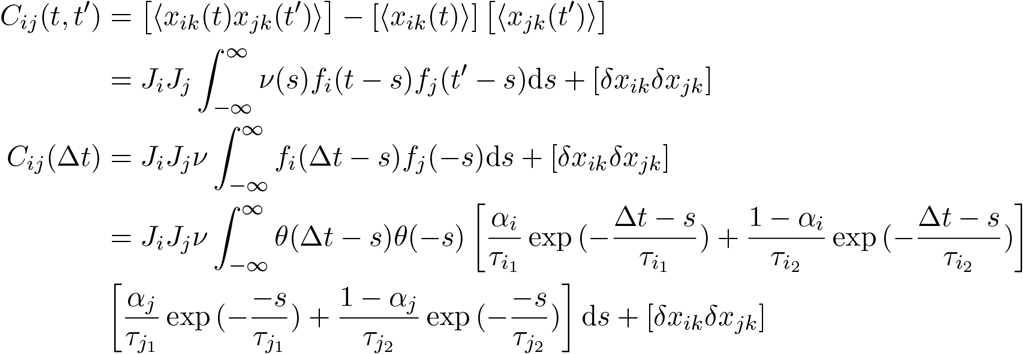

The above equation, in general (except for *i* = *j*), is asymmetric with respect to the sign of Δ*t*. For Δ*t >* 0 we have:

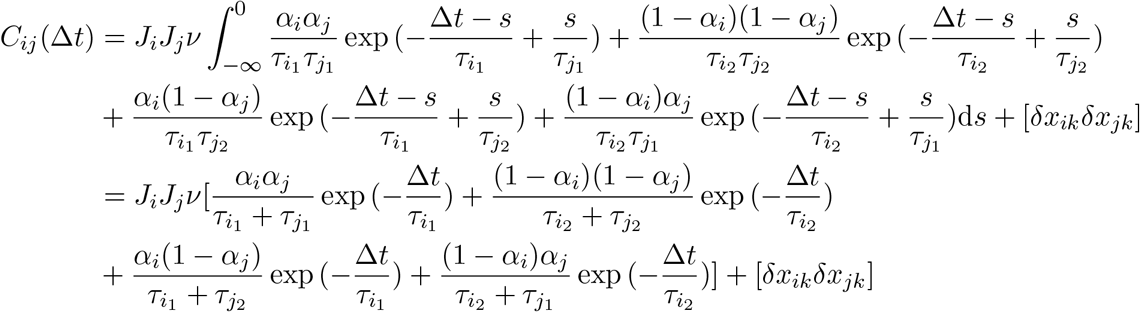

For Δ*t <* 0 we have:

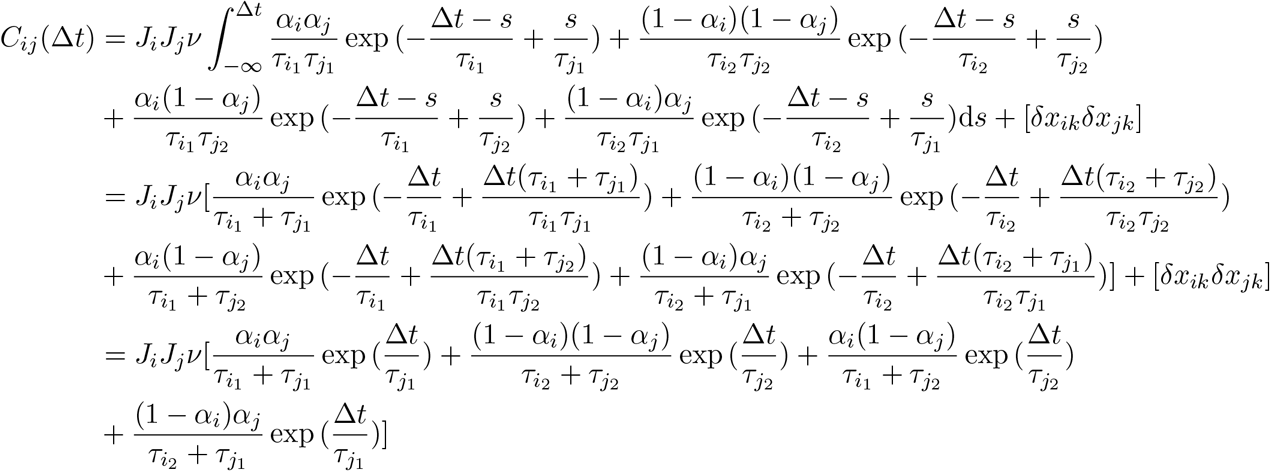

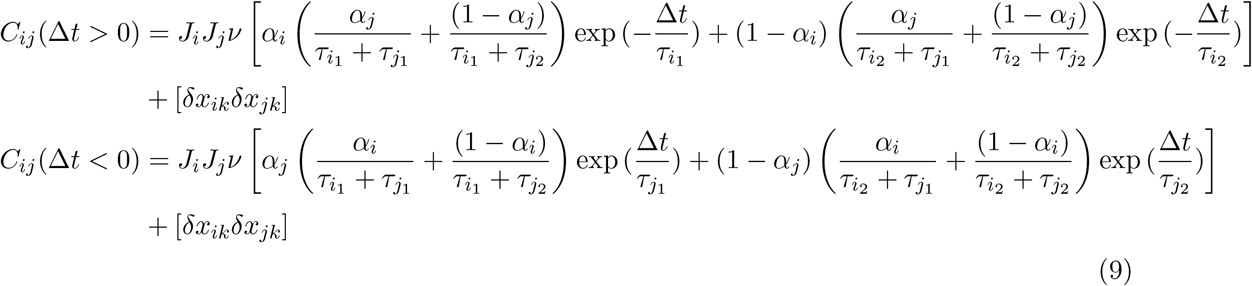

In the case *i* = *j* the correlation function becomes symmetric with respect to Δ*t*:

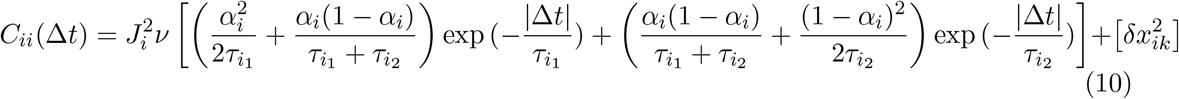

For the special case 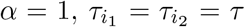, only a single decaying exponential remains, and the correlation function reduces to

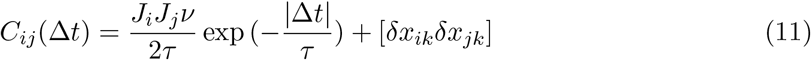

### Predicting cross-covariance amplitudes from auto-covariances

Working in normalized units with *ν/*(2*τ*) = 1, assume three event classes with fractions *e, e*_1_, *e*_2_ (*e* + *e*_1_ + *e*_2_ = 1) and binary targeting: head size (hs) is hit by *e* and *e*_1_; neck length (nl) and neck width (nw) by *e* and *e*_2_. The auto-covariance amplitudes are then

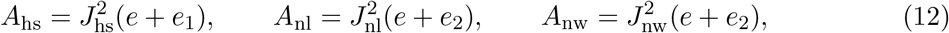

which fix the jump sizes |*J*_*i*_|.

The cross-covariance amplitudes, *A*_*ij*_ = |*J*_*i*_*J*_*j*_| × (fraction of events hitting both *i, j*), follow as

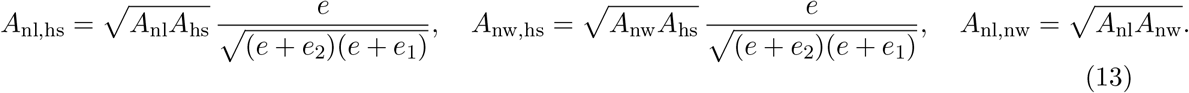

In the fully coordinated limit *e* = 1, all three reduce to the geometric mean of the corresponding auto-covariances, 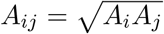.

Varying *e, e*_1_, *e*_2_ traces the predicted cross-covariance surface (Fig. 5d). The fully coordinated case (*e* = 1) and the head-or neck-dominated limits (*e*_1_ ≫ *e, e*_2_ or *e*_2_ ≫ *e, e*_1_) all fall outside the 95% credible interval of the measured cross-covariances; only an intermediate mixture of the three classes is consistent with the data, over the ranges given in the main text (Fig. 5).

### Limitations of an activity-independent stochastic model

If synaptic turnover were driven solely by the stochastic dynamics of actin filaments, morphological changes would be independent of pre- and postsynaptic activity. Previous work has shown that the complex dynamics of actin and the spine membrane can be reduced to the number of polymerized actin filaments [34], with spine area changes proportional to this number at a given time.

Approximating actin polymerization as proportional to the spine morphological measures, the dynamics of each feature follow the stochastic differential equation 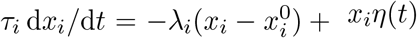, where *x*_*i*_(*t*) is the feature value, *τ*_*i*_ a timescale, *λ*_*i*_ the relaxation strength toward the preferred value 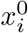, and *η*(*t*) Gaussian white noise. The multiplicative noise term reflects the stochastic nature of actin polymerization, which scales with the current spine size.

This model yields correlation functions decaying on the order of *τ* (minutes to hours), far below the multi-day timescale observed in the data (Figs. 2 and S2). It further predicts zero cross-correlation between different features, in contrast to the robust nonzero cross-correlations measured here. An activity-independent actin model therefore cannot account for the observations (Fig. 2).

### Coordinated events dominate current variability across morphospace

The counterfactual reconstructions of Fig. 6c were computed for a single spine. To confirm this does not depend on that choice, we repeated them at six locations in morphospace, each setting one feature to a near-extreme observed day-14 value while the other two start at the population mean (Fig. S5). At every location, reconstructions retaining coordinated events reproduced the broad, right-skewed current distribution of the full model.

**Figure S1:**
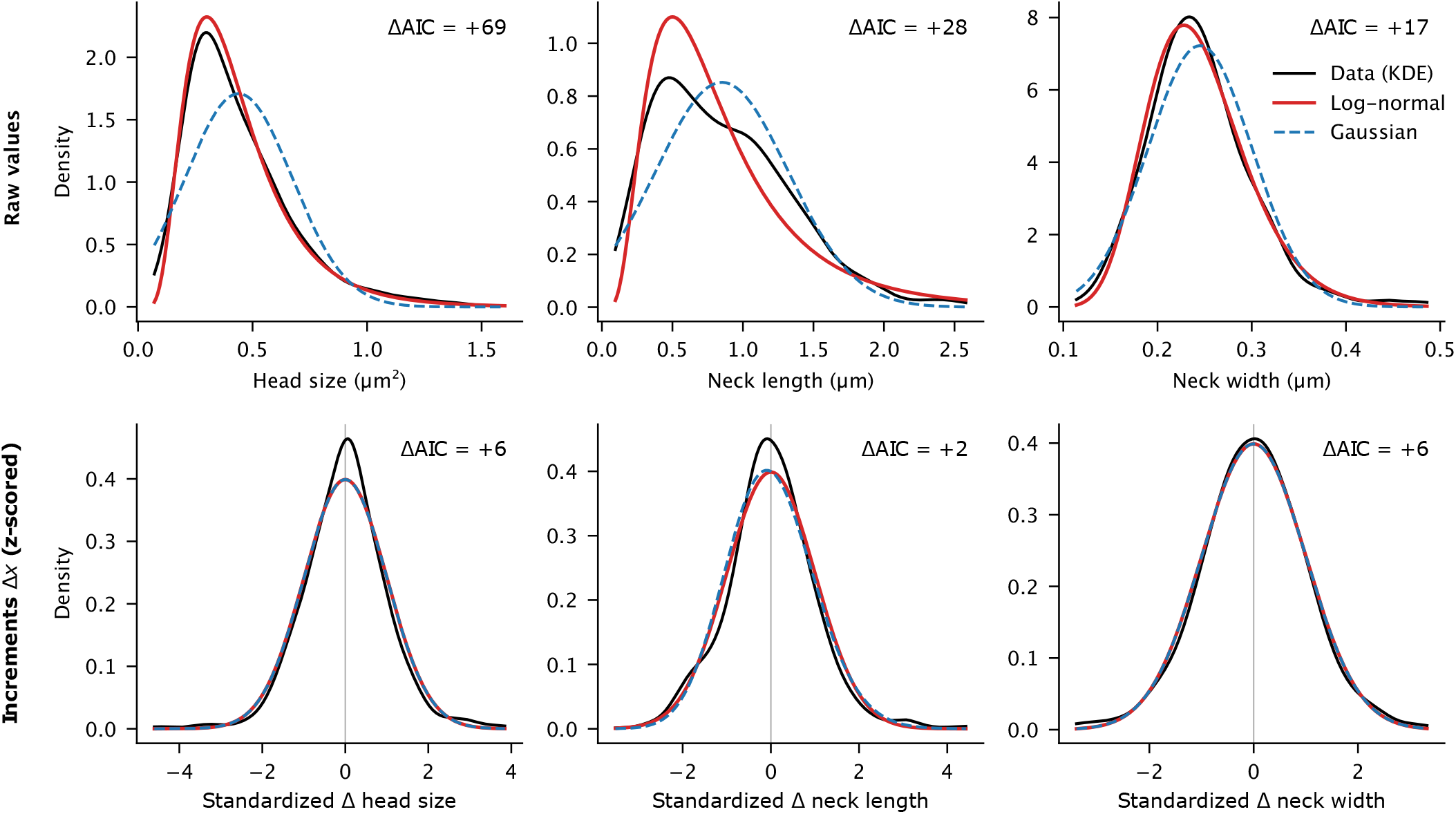
Morphological features are log-normal and their increments Gaussian. **Top:** spine-weighted distributions of the raw values of head size, neck length, and neck width (black, kernel density estimate) with log-normal (red) and Gaussian (dashed blue) fits; the log-normal is favored for all three features. **Bottom:** between-timepoint increments Δ*x*, pooled across lags after standardizing each lag by its own spine-weighted mean and standard deviation, with a standard normal *N* (0, 1) (red) and a freely fitted shifted log-normal (dashed blue); the two coincide as the log-normal converges onto the Gaussian. Each panel reports the weighted ΔAIC, defined so that positive values favor the better-supported model.

**Figure S2:**
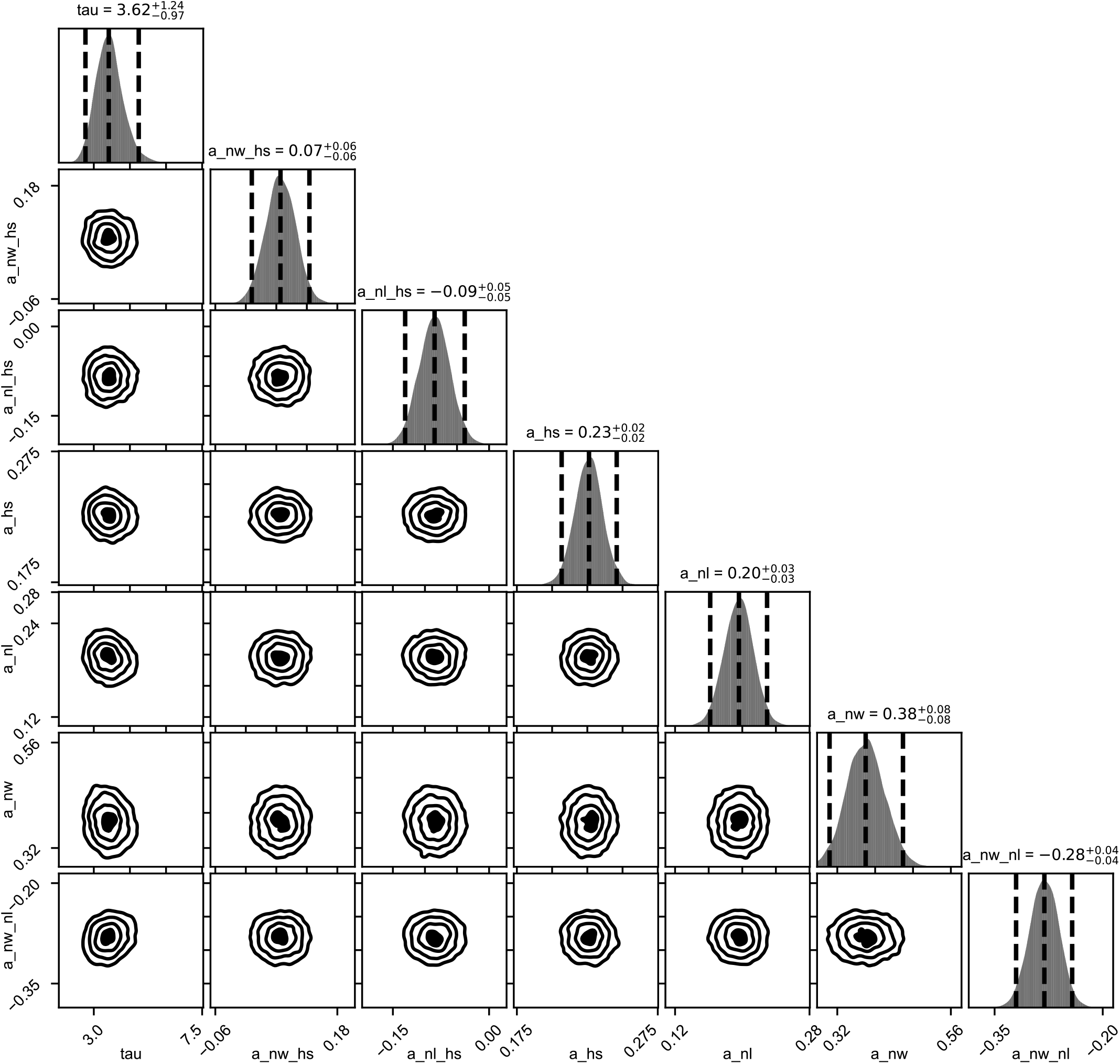
Posterior distributions for the one-timescale model. Corner plot showing posterior distributions of the single timescale *τ* and amplitude parameters for all autocorrelation and cross-correlation functions.

**Figure S3:**
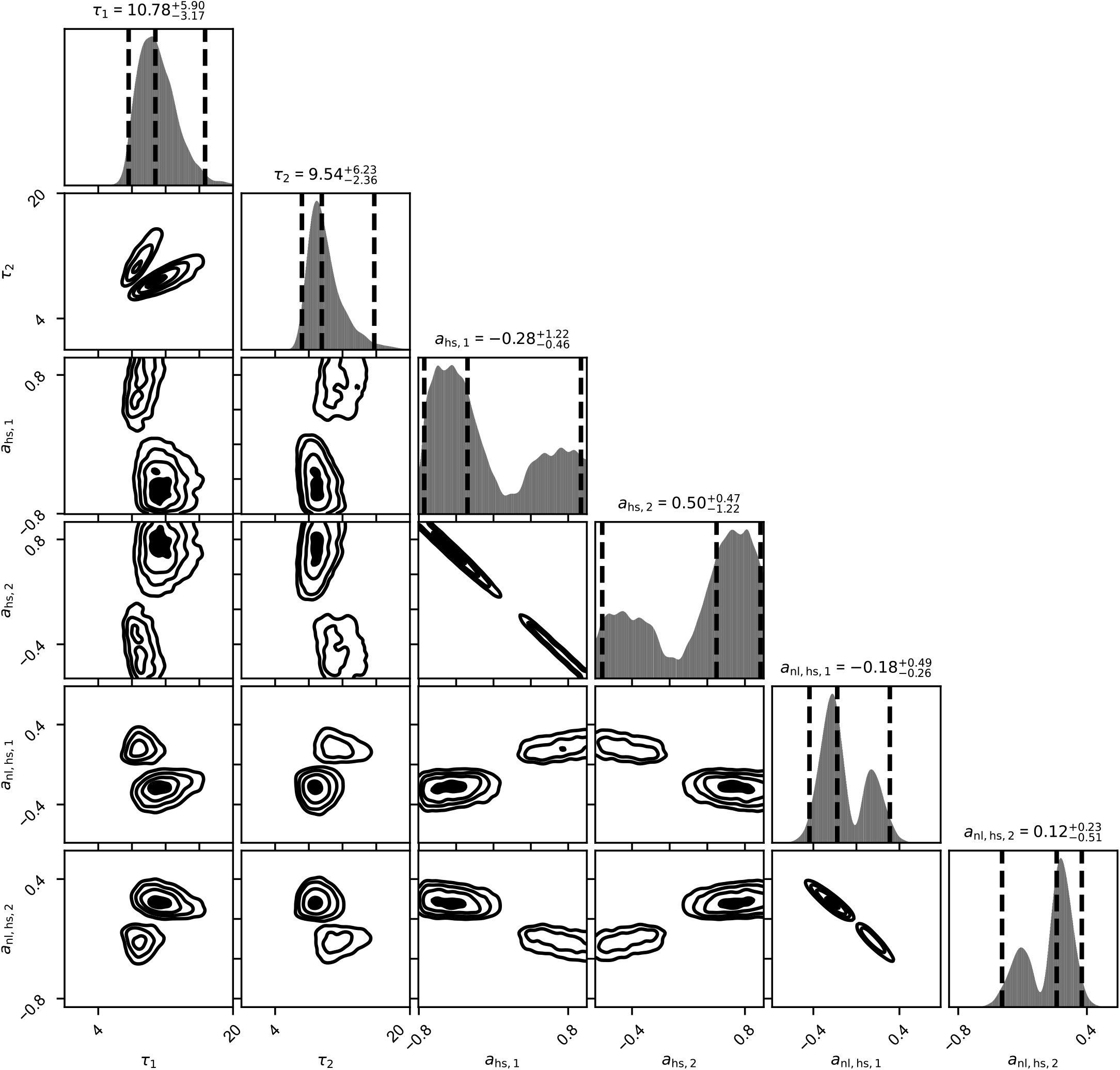
Posterior distributions for the two-timescale model. Corner plot showing the two fitted timescales (*τ*_1_, *τ*_2_) and amplitudes for head-size autocorrelation and head–neck correlation. The two timescales strongly overlap, and amplitude parameters exhibit ridges, indicating non-identifiability. The two-timescale model is therefore not supported by the data.

**Figure S4:**
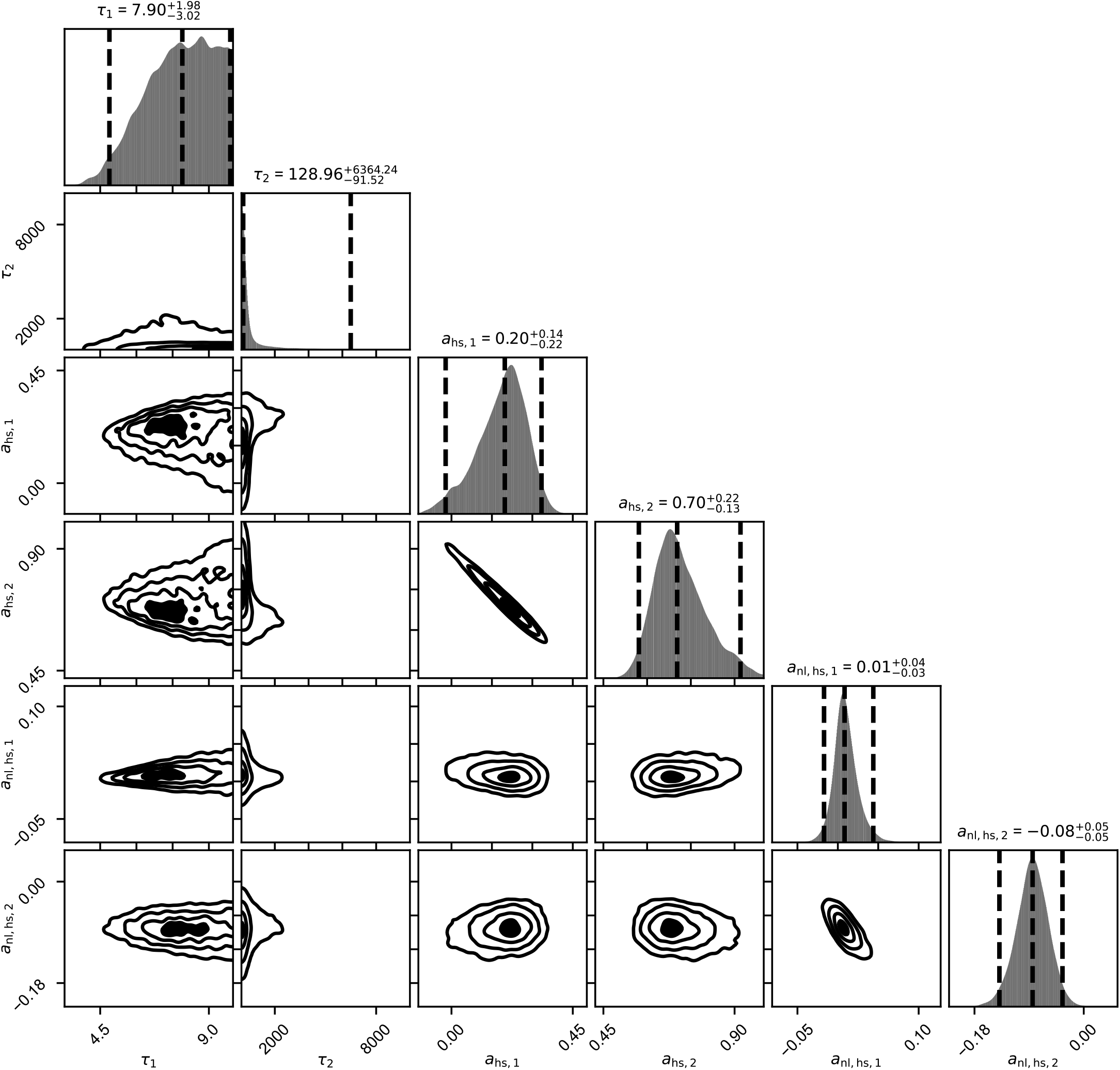
Posterior distributions for the two-timescale model without offset. Corner plot showing the two fitted timescales (*τ*_1_, *τ*_2_) and amplitudes for head-size auto-correlation and head–neck correlation. Lacking a persistent offset, the model can only reproduce the non-vanishing long-lag correlations by pushing the second timescale *τ*_2_ to enormous, unconstrained values (posterior median ~130 days with a credible interval extending to thousands of days, i.e. years), effectively mimicking a constant baseline with an arbitrarily slow exponential. The resulting parameters are poorly constrained, showing that a quenched, spine-specific offset—rather than a second slow decay—is required, and that the data are not consistent with a purely decaying two-timescale process.

**Figure S5:**
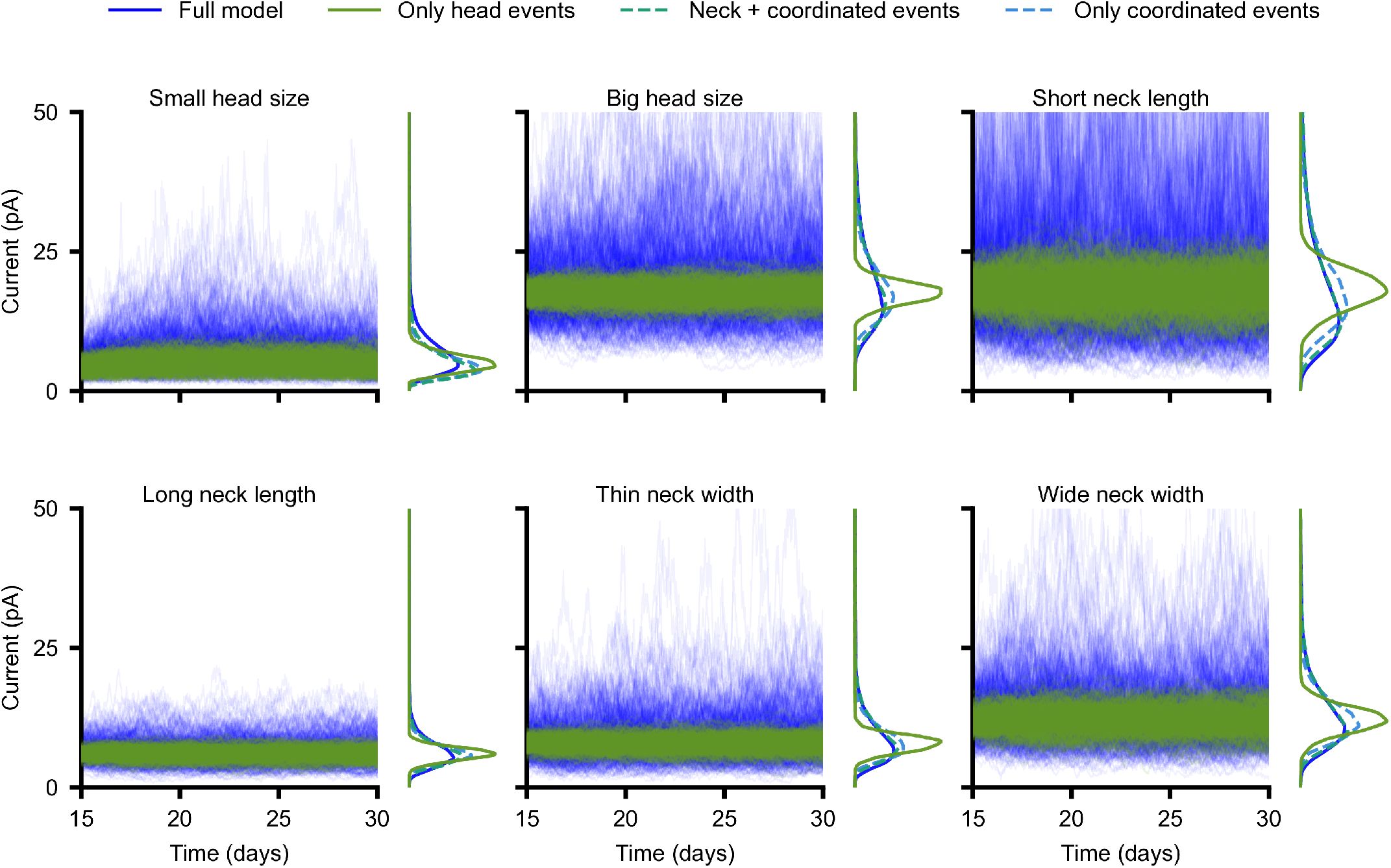
Coordinated events dominate current variability across morphospace. The counterfactual reconstructions of Fig. 6c, repeated at six morphospace locations (one feature at a near-extreme day-14 value, the other two starting at the population mean). Each panel shows predicted synaptic current (days 15–30) for the full model (blue) and reconstructions retaining only head events (orange), neck plus coordinated events (dashed blue), or only coordinated events (dashed green), with the corresponding current distributions (days 20–30) alongside. Every reconstruction containing coordinated events reproduces the broad, right-skewed distribution of the full model.

## Bibliography

[1] Functional connectomics spanning multiple areas of mouse visual cortex. Nature, 640(8058):435–447, 2025.

[2] Jaime Grutzendler, Narayanan Kasthuri, and Wen-Biao Gan. Long-term dendritic spine stability in the adult cortex. Nature, 420(6917):812–816, 2002.

[3] Anthony JGD Holtmaat, Joshua T Trachtenberg, Linda Wilbrecht, Gordon M Shepherd, Xiaoqun Zhang, Graham W Knott, and Karel Svoboda. Transient and persistent dendritic spines in the neocortex in vivo. Neuron, 45(2):279–291, 2005.

[4] Yi Zuo, Aerie Lin, Paul Chang, and Wen-Biao Gan. Development of long-term dendritic spine stability in diverse regions of cerebral cortex. Neuron, 46(2):181–189, 2005.

[5] Tim VP Bliss and Graham L Collingridge. A synaptic model of memory: long-term potentiation in the hippocampus. Nature, 361(6407):31–39, 1993.

[6] Stephen J Martin, Paul D Grimwood, and Richard GM Morris. Synaptic plasticity and memory: an evaluation of the hypothesis. Annual review of neuroscience, 23(1):649–711, 2000.

[7] Masanori Matsuzaki, Graham CR Ellis-Davies, Tomomi Nemoto, Yasushi Miyashita, Masamitsu Iino, and Haruo Kasai. Dendritic spine geometry is critical for ampa receptor expression in hippocampal ca1 pyramidal neurons. Nature neuroscience, 4(11):1086–1092, 2001.

[8] Jan Tønnesen, Gergely Katona, Balázs Rózsa, and U Valentin Nägerl. Spine neck plasticity regulates compartmentalization of synapses. Nature neuroscience, 17(5):678–685, 2014.

[9] Roberto Araya, Tim P Vogels, and Rafael Yuste. Activity-dependent dendritic spine neck changes are correlated with synaptic strength. Proceedings of the National Academy of Sciences, 111(28):E2895–E2904, 2014.

[10] David Holcman and Rafael Yuste. The new nanophysiology: regulation of ionic flow in neuronal subcompartments. Nature Reviews Neuroscience, 16(11):685–692, 2015.

[11] Jerome Cartailler, Zeev Schuss, and David Holcman. Electrostatics of non-neutral biological microdomains. Scientific reports, 7(1):11269, 2017.

[12] Jerome Cartailler, Taekyung Kwon, Rafael Yuste, and David Holcman. Deconvolution of voltage sensor time series and electro-diffusion modeling reveal the role of spine geometry in controlling synaptic strength. Neuron, 97(5):1126–1136, 2018.

[13] Ken-Ichi Okamoto, Takeharu Nagai, Atsushi Miyawaki, and Yasunori Hayashi. Rapid and persistent modulation of actin dynamics regulates postsynaptic reorganization underlying bidirectional plasticity. Nature neuroscience, 7(10):1104–1112, 2004.

[14] Naoki Honkura, Masanori Matsuzaki, Jun Noguchi, Graham CR Ellis-Davies, and Haruo Kasai. The subspine organization of actin fibers regulates the structure and plasticity of dendritic spines. Neuron, 57(5):719–729, 2008.

[15] Anthony Holtmaat and Karel Svoboda. Experience-dependent structural synaptic plasticity in the mammalian brain. Nature Reviews Neuroscience, 10(9):647–658, 2009.

[16] Gianluigi Mongillo, Simon Rumpel, and Yonatan Loewenstein. Intrinsic volatility of synaptic connections—a challenge to the synaptic trace theory of memory. Current opinion in neurobiology, 46:7–13, 2017.

[17] Ayse Dosemeci, Richard J Weinberg, Thomas S Reese, and Jung-Hwa Tao-Cheng. The postsynaptic density: there is more than meets the eye. Frontiers in synaptic neuroscience, 8:23, 2016.

[18] Harold D MacGillavry, Yu Song, Sridhar Raghavachari, and Thomas A Blanpied. Nanoscale scaffolding domains within the postsynaptic density concentrate synaptic ampa receptors. Neuron, 78(4):615–622, 2013.

[19] Ai-Hui Tang, Haiwen Chen, Tuo P Li, Sarah R Metzbower, Harold D MacGillavry, and Thomas A Blanpied. A trans-synaptic nanocolumn aligns neurotransmitter release to receptors. Nature, 536(7615):210–214, 2016.

[20] Maria Fischer, Stefanie Kaech, Darko Knutti, and Andrew Matus. Rapid actin-based plasticity in dendritic spines. Neuron, 20(5):847–854, 1998.

[21] Albrecht Sigler, Won Chan Oh, Cordelia Imig, Bekir Altas, Hiroshi Kawabe, Benjamin H Cooper, Hyung-Bae Kwon, Jeong-Seop Rhee, and Nils Brose. Formation and maintenance of functional spines in the absence of presynaptic glutamate release. Neuron, 94(2):304–311, 2017.

[22] Noam E Ziv and Naama Brenner. Synaptic tenacity or lack thereof: spontaneous remodeling of synapses. Trends in neurosciences, 41(2):89–99, 2018.

[23] U Valentin Nägerl, Katrin I Willig, Birka Hein, Stefan W Hell, and Tobias Bonhoeffer. Live-cell imaging of dendritic spines by sted microscopy. Proceedings of the National Academy of Sciences, 105(48):18982–18987, 2008.

[24] Hanna Steffens et al. Stable but not rigid: Chronic in vivo sted nanoscopy reveals extensive remodeling of spines, indicating multiple drivers of plasticity. Science Advances, 7, 2021.

[25] Waja Wegner, Heinz Steffens, Carola Gregor, Fred Wolf, and Katrin I Willig. Environmental enrichment enhances patterning and remodeling of synaptic nanoarchitecture as revealed by sted nanoscopy. Elife, 11:e73603, 2022.

[26] Katrin I Willig. In vivo super-resolution of the brain–how to visualize the hidden nanoplasticity? Iscience, 25(9), 2022.

[27] Matthew Larkum. A cellular mechanism for cortical associations: an organizing principle for the cerebral cortex. Trends in neurosciences, 36(3):141–151, 2013.

[28] Friedrich Schuessler, Francesca Mastrogiuseppe, Alexis Dubreuil, Srdjan Ostojic, and Omri Barak. The interplay between randomness and structure during learning in rnns. Advances in neural information processing systems, 33:13352–13362, 2020.

[29] Francesca Mastrogiuseppe and Srdjan Ostojic. Linking connectivity, dynamics, and computations in low-rank recurrent neural networks. Neuron, 99(3):609–623, 2018.

[30] Nicolas Brunel. Is cortical connectivity optimized for storing information? Nature neuroscience, 19(5):749–755, 2016.

[31] Johannes Schindelin, Ignacio Arganda-Carreras, Erwin Frise, Verena Kaynig, Mark Longair, Tobias Pietzsch, Stephan Preibisch, Curtis Rueden, Stephan Saalfeld, Benjamin Schmid, et al. Fiji: an open-source platform for biological-image analysis. Nature methods, 9(7):676–682, 2012.

[32] Joshua S Speagle. dynesty: a dynamic nested sampling package for estimating bayesian posteriors and evidences. Monthly Notices of the Royal Astronomical Society, 493(3):3132–3158, 2020.

[33] Kenneth P. Burnham and David R. Anderson. Model Selection and Multimodel Inference: A Practical Information-Theoretic Approach. Springer, New York, 2nd edition, 2002.

[34] Mayte Bonilla-Quintana, Florentin Wörgötter, Christian Tetzlaff, and Michael Fauth. Modeling the shape of synaptic spines by their actin dynamics. Frontiers in synaptic neuroscience, 12:9, 2020.

